# Clonally expanded IgG antibody-secreting cells preferentially target influenza nucleoprotein following homologous and heterologous infections

**DOI:** 10.1101/2024.12.09.627526

**Authors:** Andreas Agrafiotis, Raphael Kuhn, Camilla Panetti, Marco Venerito, Hathaichanok Phandee, Lucas Stalder, Danielle Shlesinger, Vittoria Martinolli D’Arcy, Kai-Lin Hong, Daphne van Ginneken, Alessandro Genovese, Nicole Joller, Annette Oxenius, Sai T. Reddy, Alexander Yermanos

## Abstract

Infection with influenza virus remains a significant global health concern due to its ability to acquire mutations at key antigenic sites to escape antibody recognition. While germinal center (GC) and memory B cells have been well studied following influenza infection, the clonal dynamics of antibody secreting cells (ASCs), particularly those within the bone marrow (BM) niche that are responsible for serum immune protection, remain poorly understood. Here, we combine single-cell RNA (scRNA) and B cell receptor (BCR) sequencing to characterize individual ASCs following various Influenza exposure histories. We find that BM repertories are populated by highly expanded and class-switched ASCs following Influenza infection with similar transcriptional and repertoire characteristics regardless of homologous or heterologous infection histories. By combining single-cell analysis with monoclonal antibody expression and characterization, we find that a large proportion of the expanded IgG-, but not IgA-, ASC repertoire demonstrates specificity to influenza nucleoprotein (NP). Together, our data reveal the complex relationship between BM ASC repertoires, mucosal humoral immune responses, and BCR antigen specificity during influenza infection.

## Introduction

Influenza virus is a common airborne pathogen that infects cells of the respiratory tract and represents a major threat to human health (Krammer et al., 2018). Antibodies targeting the head domain of the viral hemagglutinin (HA), a surface glycoprotein essential for viral entry into host cells, typically exhibit potent neutralizing activity and have been identified as a correlate of protection (Altman et al., 2018). However, in contrast to the relatively stable genes of influenza that encode internal proteins like NP, the genes for viral glycoproteins such as HA and neuraminidase (NA) accumulate amino acid substitutions over time. These changes, driven by the selective pressure exerted by population immunity, progressively alter the antigenic properties of the glycoproteins. This, combined with reassortant strains, which emerge through the exchange of genetic material between distinct viral strains, has substantially contributed to influenza’s ability to evade immune responses and complicate control efforts (Krammer, 2019).

In mice, intranasal (i.n.) infection with influenza initiates B cell responses in several organs, characterized by persistent GC formation (Adachi et al., 2015; Onodera et al., 2012). Within GCs, B cells undergo somatic hypermutation (SHM) and affinity maturation of BCRs. This process can give rise to low- and high-affinity B cell clones that contribute to both the memory B cells and the ASC compartment (plasma cells (PCs) and plasmablasts). Upon secondary infection, the activation of these cells can result in the rapid secretion and delivery of antibodies directly to the sites of viral replication in a highly localized manner (Ise et al., 2018; Rothaeusler & Baumgarth, 2010; Sprumont et al., 2023). After initial differentiation in secondary lymphoid structures or inflamed tissues, B cells can enter survival niches, such as the BM, where they can provide long-lasting protection (Benet et al., 2021; Koutsakos et al., 2019; Tellier & Nutt, 2022; Wu et al., 2016). In contrast to GC biology and memory B cell dynamics (Andrews et al., 2019; de Carvalho et al., 2023; Mathew et al., 2021; Matsuda et al., 2019; Turner et al., 2020; Victora & Wilson, 2015; W. T. Yewdell et al., 2021), much less is known regarding the functional characteristics of the BM ASC compartment following influenza infection; this is despite their importance in maintaining protective antibodies in serum (Brynjolfsson et al., 2017; Halliley et al., 2015; Robinson et al., 2022).

Single-cell sequencing (scSeq) of antibody repertoires is able to identify natively paired heavy and light chain sequences, which can be combined with recombinant antibody expression and functional (antigen-specificity) characterization. Previous studies on vaccine-induced ASC repertoires (murine- and BM-derived) demonstrated that they are dominated by highly expanded antigen-specific clones (Reddy et al., 2010; Wang et al., 2016). We have previously demonstrated in mice that a portion of the expanded IgG ASC repertoire in BM is antigen-specific following various immune challenges (Agrafiotis et al., 2023; Neumeier et al., 2021, 2022), whereas expanded IgM ASC repertoires in BM were not immunogen-specific (Neumeier et al., 2021). More recently, an unmutated IgM ASC subset has been described in naive mice that is enriched with public antibody sequences recognizing self-antigens and commensal microbes (Liu et al., 2023).

The complexity of ASCs, including their diverse origins, isotypes, longevity, and specificities, present numerous unanswered questions in the field (Robinson et al., 2022). For example, it remains underexplored how factors such as pre-existing immunity and repeated vaccination influence the following: (i) composition of the BM ASC repertoire, (ii) ASC turnover and (iii) future immune responses. In this context, immune responses to influenza are influenced by previous exposure through infection or vaccination, leading to a phenomenon known as immune imprinting. This imprinting can bias the immune system toward recognizing and responding to prior viral variants, thereby limiting the effectiveness of immune responses against newly emerging variants (Altman et al., 2018; Arevalo et al., 2020; Fonville et al., 2014; Kim et al., 2009; Oidtman et al., 2021; J. W. Yewdell & Santos, 2021). More recently, BCR fate mapping experiments have demonstrated that GCs resulting from secondary exposure are populated by naive rather than memory B cells, suggesting that immune imprinting might be a phenomenon exclusively observed on the serum level (Mesin et al., 2020; Schiepers et al., 2023). In influenza infection, immune imprinting is often associated with antibodies targeting the immunodominant HA, with previous studies showcasing antibody backboosting in serum (Arevalo et al., 2020; Linderman & Hensley, 2016). How serum-antibody backboosting is reflected on a single-cell level in ASCs still remains undetermined.

In this study, we used scSeq and computational analysis of murine antibody repertoires combined with antibody-antigen-binding measurements to explore the functional characteristics of individual ASCs generated from different influenza infection immune histories. We focused on the BM compartment given its crucial role in accommodating terminally differentiated ASCs, which reside for extended periods following infection or vaccination and are essential for long-term humoral immunity and sustained antibody production (Manz et al., 2005; Turner et al., 2021). Our findings reveal expanded ASC repertories following influenza infection marked by abundant levels of highly expanded IgA and lower levels of expanded IgG clones. Intriguingly, infection history appears to have a limited impact on the repertoire diversity and transcriptional profiles of ASC repertoires. Characterizing the biophysical properties of highly expanded ASCs demonstrated frequent IgG and rare IgA specificity to influenza NP rather than the more commonly expected targeted surface glycoproteins (HA and NA). In addition, a fraction of the expanded IgA clones showed signs of polyreactivity against self and non-self antigens. Our results highlight the complex relation between ASC repertoires, mucosal-associated humoral immune responses and antibody-antigen specificity during influenza infection.

## Results

### Highly expanded and class-switched ASC clones are found in the BM following influenza infection

To investigate how influenza infection history influences the function and selection of ASCs, we performed scSeq of murine BM following homologous and heterologous intranasal (i.n.) infections from multiple batches (Figure 1A). More specifically, PR8 (H1N1) was used for primary and secondary challenge in homologous infections (PP group). For heterologous infections, it was important to choose a strain that would not be neutralized by antibodies specific for the HA or NA glycoproteins of the priming virus (Chen et al., 2004). For this purpose, X31, a reassortant strain with H3N2 glycoprotein gene segments (HA and NA) and the remaining genes from PR8, was used (PX group). ASCs (CD138hi) were isolated by fluorescence-activated cell sorting (FACS) at 25 days post-secondary infection and serum titers were assessed using enzyme-linked immunosorbent assay (ELISA) against influenza surface proteins HA and neuraminidase (NA) and the internal nucleoprotein (NP) (Figure 1B). Detectable titers were observed for all three antigens and minimal cross-reactivity was observed between HA1 (PR8) and HA3 (X31) glycoproteins in homologously infected samples (Figure 1B). Following B cell isolation, we performed scSeq of transcriptomes and antibody repertories using the 5’ GEX and V(D)J protocols from 10x Genomics. Subsequent to scSeq and quality filtering of cells, uniform manifold approximation projection (UMAP) and unsupervised clustering were performed for an overview of gene expression for all cell populations (Figure 1C). Visualizing the expression of known B cell markers demonstrated that several clusters were characterized by an ASC-like phenotype (e.g. *Cd138 (Sdc1), Xbp1, Slamf7, Slpi),* while others almost completely lacked expression of such markers and exhibited high expression of early development and mature B cell genes like *B220* (*Ptprc*), *Cd19* and *Pax5* (Figure 1D). This may partially reflect the limitations in our sorting methodology and time constraints during sample processing, potentially leading to the inclusion of non-ASC B cell populations in our dataset. To further characterize B cell phenotypic diversity, we performed differential gene expression analysis and identified distinct B cell modules based on expression of known B cell markers (Figure 1E, Figure S1) (Mathew et al., 2020). This highlighted a clear separation between naive-like B cells and ASCs, with the latter found primarily in clusters 1, 2, 3, 4, 7, 9 and 12. Interestingly, cluster 7 was also characterized by expression of *Mki67*, indicating the presence of actively proliferating ASCs (Figure 1D, Figure S2).

**Figure 1.**
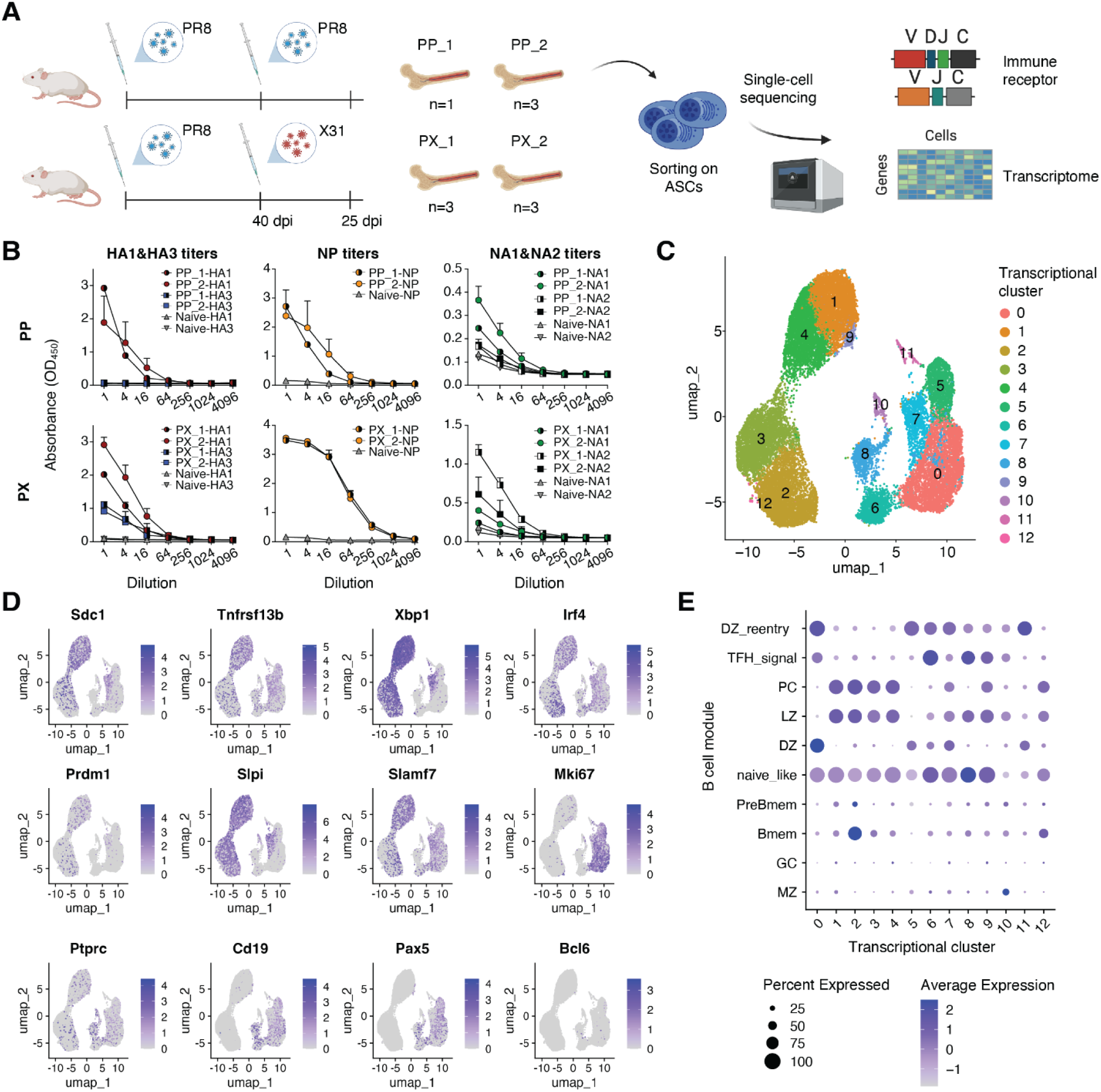
Single-cell sequencing of transcriptomes and antibody repertoires following influenza infection. A) Experimental overview of influenza infection, B cell isolation and single-cell sequencing of transcriptomes and antibody repertoires (dpi = days post infection). B) Antibody titers per condition measured by ELISA with four-fold pre-diluted serum (1:100) from uninfected mouse (naive) and influenza infected mice (PR8PR8 (PP) and PR8X31 (PX)) tested against hemagglutinin (HA) (left), neuraminidase (NA) (right) and nucleoprotein (NP) (middle). Legend indicates condition_batch-capture protein. C) Uniform manifold approximation projection (UMAP)-based total gene expression for B cells following PP and PX infections. D) UMAP-based gene expression of generic B cell markers. E) Dottile plot showing B cell subset assignment across all clusters based on expression of genes defining different B cell phenotypes. The intensity of each dot corresponds to the average expression of all cells within a given cluster, and the size corresponds to the percentage of cells with detectable gene expression.

Next, we leveraged the capabilities of scSeq to integrate transcriptome information with antibody repertoire features. To this end, cells having the same fully rearranged common ancestor were grouped together using enclone program within cellranger software (10X Genomics), which defines clones on criteria such as identical V-J genes, matching complementarity-determining region 3 (CDR3) segment lengths, shared SHM outside the junction regions, and a minimum CDR3 nucleotide (nt) identity of at least 85% (Zheng et al., 2017). We initially visualized the fraction of B cell clones that were supported by two or more unique cells (referred to as expanded clones) (Figure 2A). This indicated that clonal expansion was primarily derived from ASC-like clusters (Figure 2A, 2B). Clonal expansion further correlated with isotype distribution, as most of the expanded cells were present in clusters with high levels of IgA and IgG expression (Figure 2B). Upon closer inspection of the 50 most expanded clones in each group, we observed that the majority of highly expanded clones were IgA-dominated across multiple independent experiments (Figure 2C). Furthermore, we observed a strong correlation between class-switched IgA clones and the number of amino acid (aa) sequence variants existing within an individual clone, which was particularly apparent within the CDR3 of the variable heavy chain (CDRH3) (Figure 2D). ASCs from mice receiving single infections with PR8 and X31 strains at 15 days post-infection exhibited similar class-switching and expansion levels compared to re-infected mice (Figure S2).

**Figure 2.**
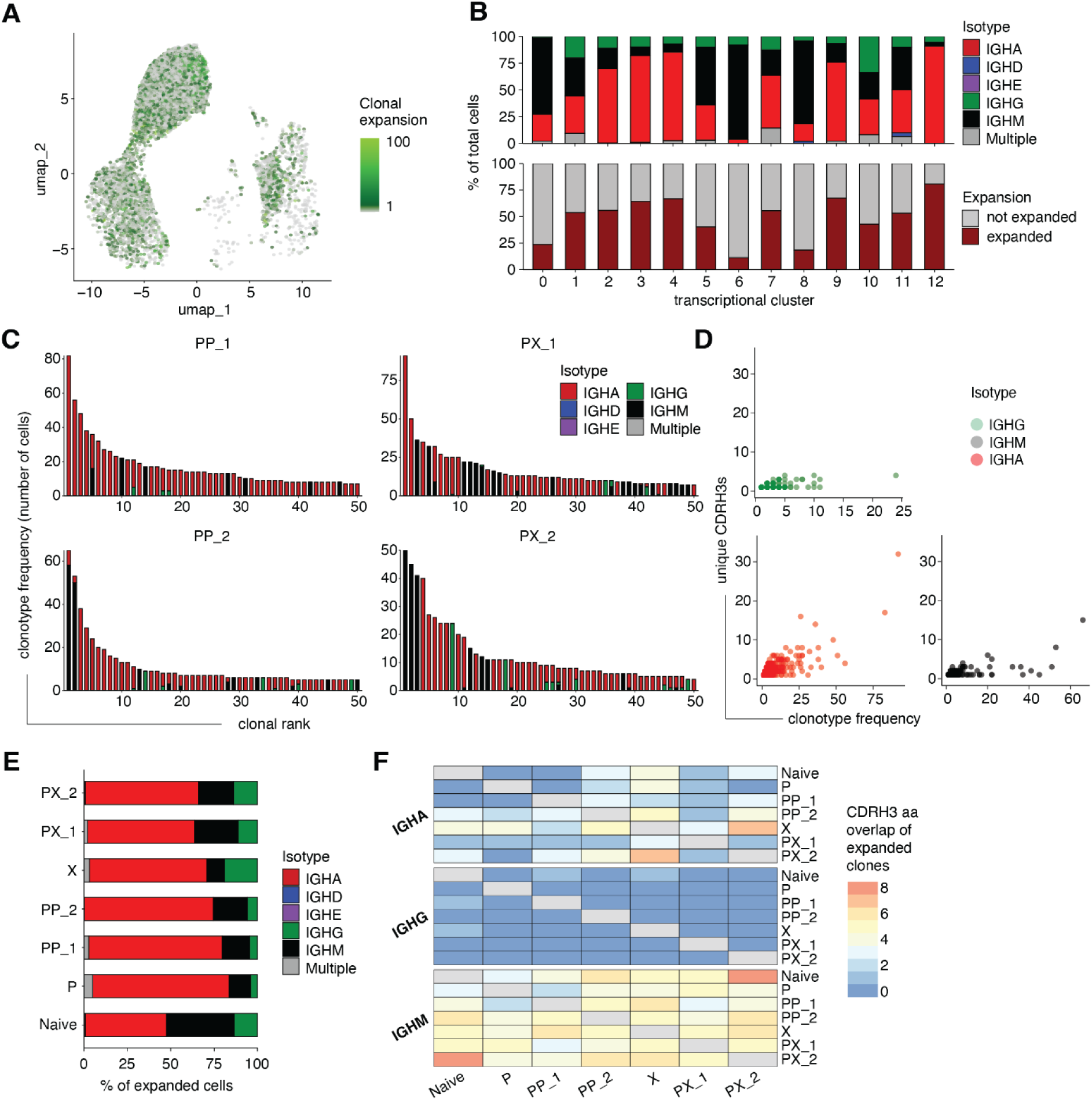
BM repertoires are enriched with expanded and IgA-expressing ASC clones. A) UMAP depicting clonal expansion for single-cells with transcriptional and antibody repertoire overlap. Values have been normalized to a scale from 1 to 100, where 1 represents a cell belonging to an unexpanded clone (gray color) and 100 to the maximum expansion (green). B) Clonal isotype and expansion distribution across transcriptional clusters. C) Clonal expansion profiles for the 50 most expanded clones colored by isotope for each condition and batch. D) Correlation between the number of unique CDRH3 aa sequence variants and the number of cells per clone. E) Isotype distribution of expanded cells across conditions. F) CDRH3 aa sequence overlap of expanded cells per isotype across conditions.

We next questioned how these findings compare to the ASC repertoire of uninfected mice and housed in specific-pathogen free (SPF) conditions. To this end we performed scSeq of ASCs from BM of uninfected mice and observed a relative enrichment of highly mutated class-switched IgA cells within the most expanded ASC clones (Figure S3). We next quantified the isotype distribution of expanded ASC clones across different conditions and batches. This showed that the expanded naive B cell repertoires exhibited the highest proportion of IgM-expressing cells with lower levels of expanded IgA ASCs compared to infected samples (Figure 2E). To determine whether ASC clones identified in infected mice were also present in naive mice, we examined CDRH3 aa sequence overlap across repertoires (Figure 2F). This revealed that IgM CDRH3 sequences were the most frequently shared across groups. This increased overlap of IgM sequences may be related to previously identified public clones, which are shared among different individual mice and found to be enriched in ASC populations (Liu et al., 2023). Interestingly, while IgM clonal overlap remained similar on the nucleotide level, the number of IgA CDRH3 nucleotide shared sequences decreased (Figure S3F). This overlap among IgA clones observed here is unlikely to result from convergent V(D)J recombination, but may instead arise through mechanisms driven by affinity maturation.

### Minor differences in transcriptional and repertoire features of ASCs are observed between homologous and heterologous influenza infection

We subsequently restricted our analysis to cells corresponding to the ASC phenotype and repeated quality filtering of cells, uniform manifold approximation projection (UMAP) and unsupervised clustering (Figure 3A, Figure S4). To understand how immune imprinting associated antibody responses in serum are reflected at the single-cell level, we next investigated whether influenza repertoire and transcriptional characteristics of the ASC compartment would be different between homologous and heterologous infections. Upon initial visualization using UMAP, we observed minor differences between ASCs isolated from mice with heterologous- or homologous-infection (Figure 3A, 3B). However, some transcriptional clusters showed batch dependency (Figure S4B). We next performed differential gene expression analysis stratified by batch (Figure S5A, S5B). While these analyses did reveal a number of significantly upregulated and downregulated genes per condition, these results were not consistent across batches.

**Figure 3.**
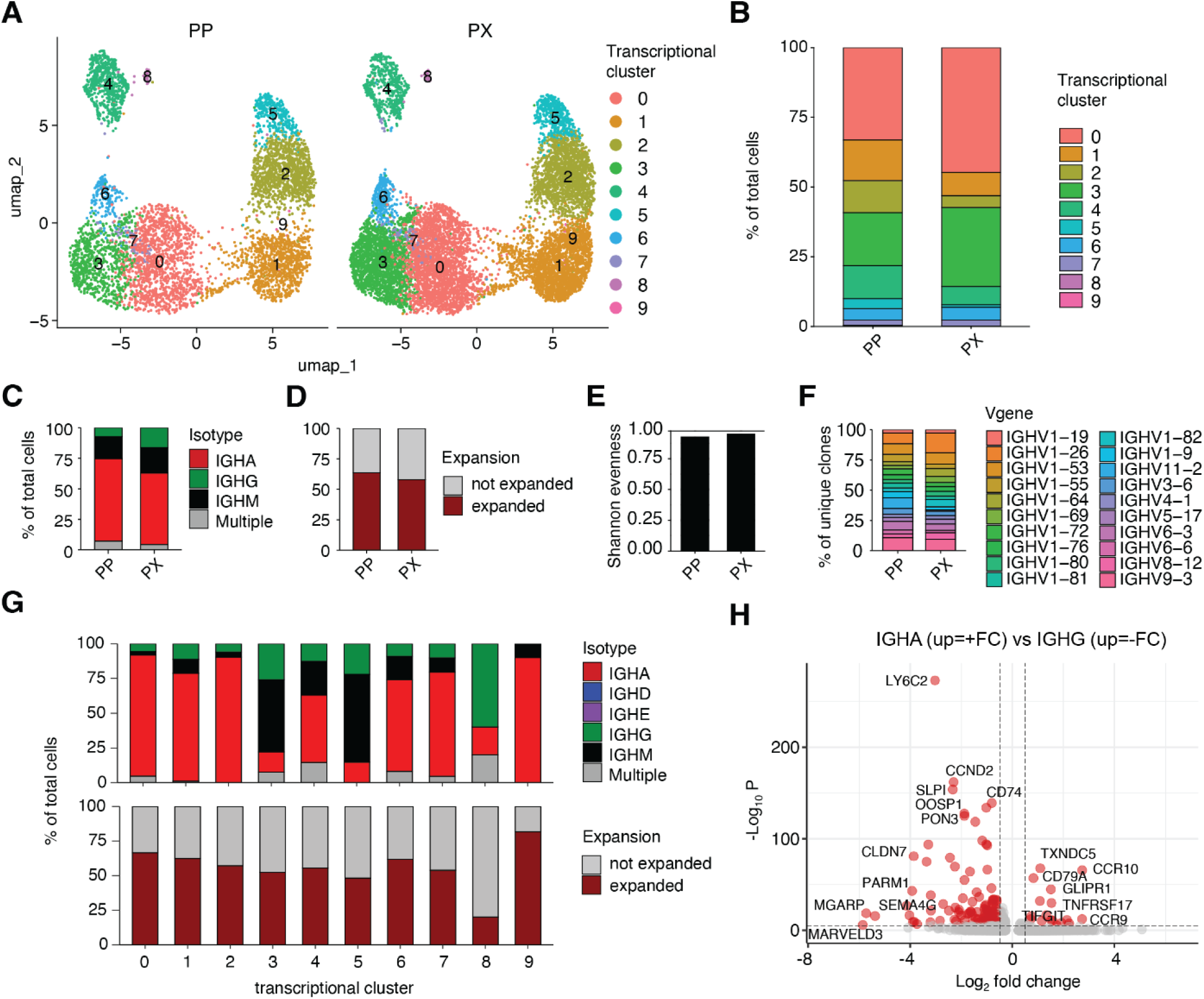
ASCs expressing different isotypes exhibit distinct transcriptional profiles independent of infection history. A) UMAP-based total gene expression of ASCs split by infection history (PR8PR8 (PP) and PR8X31 (PX)). B) Transcriptional cluster distribution in ASCs derived from mice infected with PP and PX. C-F) Repertoire feature comparisons (Isotype distribution (C), Clonal expansion (D), Shannon evenness (E) and V gene usage (F)) across conditions. G) Isotype and clonal expansion distribution across transcriptional clusters. H) Differentially expressed genes between IgA and IgG ASCs. Red dots indicate differentially expressed genes (adjusted P value < 0.01 and average log2 fold change (FC) > 0.5).

We next quantified isotype distribution and observed a minor upregulation of IgA-expressing cells in the PP group and an increased proportion of IgG-expressing cells in the PX group, in accordance with prior results (Figure 3C, Figure S5C, Figure S5D). Repertoires from both homologous- and heterologous-infected mice exhibited similar percentages of expanded clones (59% and 57%, respectively), and similar repertoire diversity and clonality, as quantified by Shannon evenness (Figure 3D, 3E, S5C, S5D). Quantification of heavy chain variable gene usage showed that no particular V gene was exclusively associated with infection history, while both groups demonstrated higher usage of the IGHV1-26, IGHV1-53, and IGHV9-53 genes (Figure 3F, Figure S5C, S5D). Finally, we quantified the mean number of SHMs per clone in the full-length variable heavy (VH) and light (VL) chain regions, which resulted in similar SHM levels for both groups (Figure S5C, S5D). Most of these findings were also validated in an independent batch that combined mice with four different influenza infection histories (Pr8Pr8, Pr8X31, X31X31, X31Pr8) (Figure S6, S7, S8). Notably, isotype distribution revealed a separation among IgG-, IgM-, and IgA-expressing cells, with IgA-expressing clusters also exhibited high levels of clonal expansion (Figure 3G, S9A). Differential gene expression analysis identified distinct genes characterizing IgG (e.g., *Ly6c2*, *Slpi*, and *Ccnd2*) and IgA (e.g., *Ccr10*, *Glpr1*, and *Cd79a*), some of which are consistent with previous findings (Agrafiotis et al., 2023; Neumeier et al., 2022; Price et al., 2019) (Figure 3H, Figure S9B). Furthermore, we observed that on average, IgA expressing cells contained more SHMs compared to IgG and IgM (Figure S9C). As Ig variable and constant genes were excluded prior to gene expression analysis, these isotype-specific profiles suggest a significant influence of isotype fate on the expression profile of ASCs. Conversely, the observed transcriptional profile differences may contribute to class switch recombination or arise from the same signals that drive switching to the respective isotype. Overall, our findings indicate minimal transcriptional and repertoire differences of ASCs arising from distinct influenza infection histories at 25 days post-infection.

### Expanded IgG clones often target NP, while IgA clones exhibit rare NP specificity and signs of polyreactivity

After characterizing the repertoire and transcriptional properties of clonally expanded ASCs arising from different infection histories, we investigated the binding profiles of their antibodies. To this end, we focused primarily on class-switched clones across different conditions that exhibited high levels of clonal expansion and SHMs. To discover the antigenic targets of these clones, we recombinantly expressed their antibodies using a murine IgG2c constant region, and interrogated antigen-specificity against different influenza viral proteins (HA, NA and NP) as well as infected cell lysate (Figure 4A). Within the IgG compartment, about 40% of expressed clones displayed clear reactivity against the viral NP (Figure 4A). While binding to NP correlated with binding to PR8 infected cell lysate, for some antibodies this correlation was less clear, which may be due to differences in the NP epitopes exposed on infected cells versus recombinant NP. Unexpectedly, the vast majority of highly expanded IgA clones failed to bind viral targets, with the exception of three lowly expanded clones, representing less than 5% of recombinantly expressed IgAs (Figure 4A, Figure S1B, Figure S4B, Figure S9B). Next, to ensure that the IgG2c constant antibody region of the expressed antibodies did not influence antigen binding, we expressed three antibodies with known specificity in both IgG and IgA backbones, which confirmed comparable binding to their cognate antigens regardless of constant region (Figure 4B). We next questioned if the observed viral reactivity in serum was exclusively due to IgG antibodies. Isotype-specific serum titers against HA1 and NP demonstrated high titers for anti-viral IgG against both antigens. IgA serum antibodies reacted above background levels against HA1, but failed to bind to NP (Figure 4C). Overall, infected mice displayed higher IgG antibody serum titers compared to naive controls, while IgA titers were comparable to naive titers (Figure 4C).

**Figure 4:**
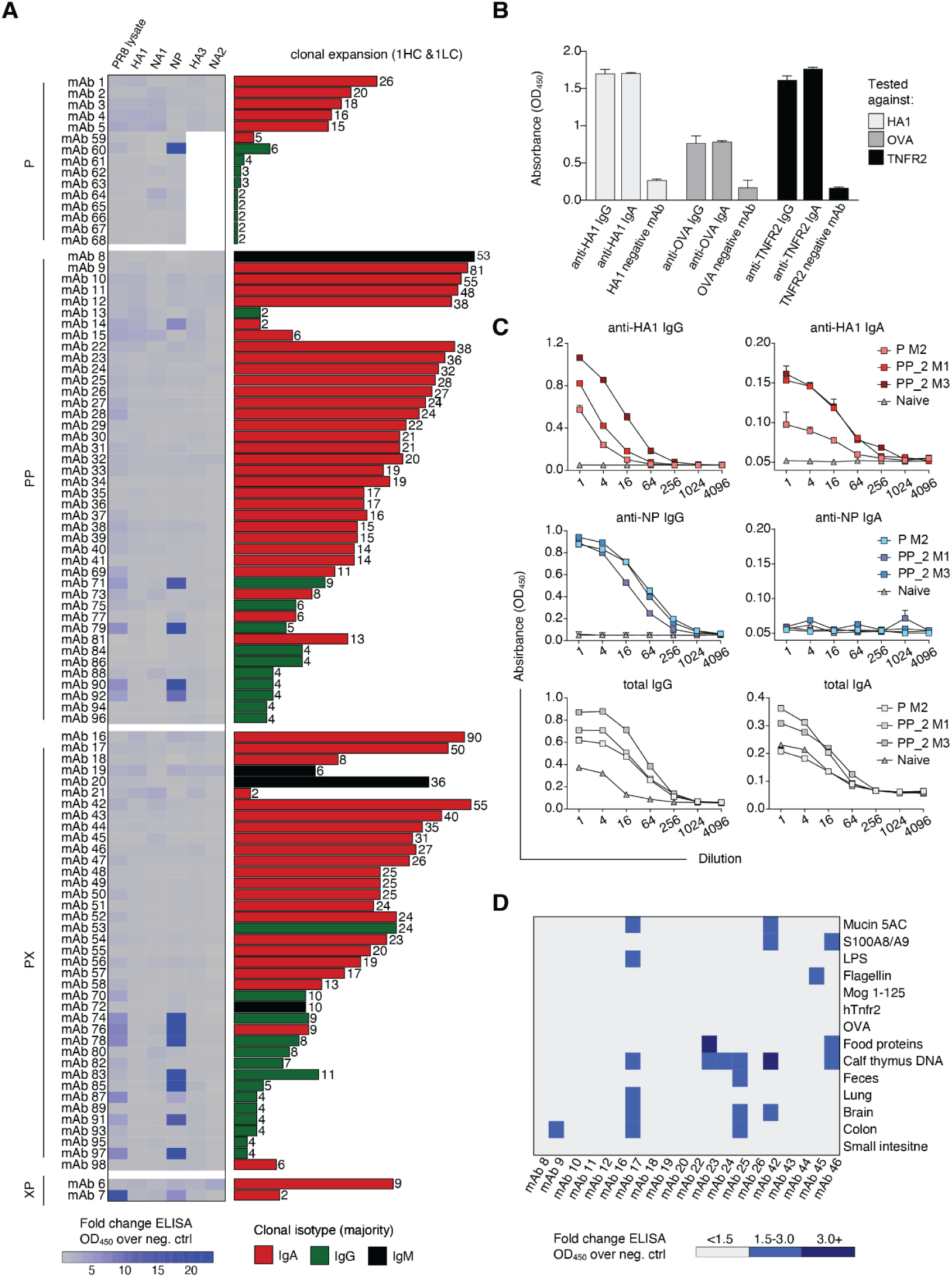
BCR specificity analysis reveals frequent NP binding within the IgG expanded ASC pool and rare NP binding of expanded IgA clones. A) Viral-protein reactivity ELISA signal values of duplicate measurements at 450 nm for individual mAbs relative to a negative background control. White color indicates mAb-antigen pairs not tested. For each mAb, the condition (left), clonal expansion and clonal isotype (right) are reported. The height of each clonal expansion bar corresponds to its normalized expansion value based on its 10x capture (see methods). The number on top of each bar indicates the number of cells with 1 heavy chain and 1 light chain. Bars are colored based on the majority isotope of that clone. More information regarding the antibody sequence can be found in the supplementary data. B) ELISA signal values of duplicate measurements at 450 nm for IgG and IgA mAbs against their cognate antigens. Legend indicates condition_batch_mouse. C) Representative serum titers of total (bottom), anti-NP (middle) and anti-HA1 (top) IGHG (left) and IGHA (right) antibodies. D) Polyreactivity ELISA signal values of duplicate measurements at 450 nm for individual mAbs relative to a negative background control.

These observations prompted us to investigate the source of IgA selection and expansion in the ASC compartment. It has previously been suggested that the mucosal compartment can contribute to the long-lived ASC pool in the BM (Liu et al., 2023; Wilmore et al., 2021). Additionally, most IgA+ ASCs in the lamina propria are GC-derived under homeostatic conditions, with substantial SHM loads (Benckert et al., 2011; Mei et al., 2009). As antibody polyreactivity and self-reactivity can be a by-product of SHMs (Meffre & Wardemann, 2008), we screened 20 antibodies originating from highly expanded IgA clones of the different infection cohorts for binding to other potential antigen sources including dietary antigens (food proteins), autoantigens (mucin 5AC, S100A8/A9, myelin oligodendrocyte glycoprotein peptide (Mog1-125), calf thymus DNA and tissue extracts), fecal proteins, microbial antigens (Lipopolysaccharide (LPS) and Flagellin) and other non-self antigens (human tumor necrosis factor receptor 2 (hTNFR2), Ovalbumin (OVA)) (Figure 4D). We observed four antibodies with potential polyreactive behavior (binding to two or more different antigens), two of which showed strong reactivity to food protein extract and calf thymus DNA (mAb 23 and mAb 42, respectively). In summary, while IgA clones showed minimal viral reactivity, a fraction of IgA-derived antibodies demonstrated binding to multiple targets.

## Discussion

Here, we set out to investigate how immune imprinting influences ASC selection and function in the context of influenza infection. We leveraged an integrated scSeq approach allowing us to recover transcriptomes from thousands of ASCs along with their naturally paired antibody VH and VL sequences and corresponding Ig isotypes. Overall, we observed highly expanded and mutated IgA ASCs overrepresented in the BM repertoire, while IgG ASCs demonstrated relatively lower expansion levels. ASCs of uninfected mice from the same animal facility also exhibited high levels of expanded and mutated IgA clones, although to a lesser extent than mice infected with influenza. Moreover, the repertoire and transcriptional features of influenza-induced ASCs were minimally influenced following heterologous and homologous infections. We have previously observed that expanded ASCs target convergent epitopes despite encoding divergent BCR sequences in the context of murine immunizations (Agrafiotis et al., 2023; Neumeier et al., 2021). Here, to elucidate the biophysical properties of influenza derived ASCs in the context of viral infection, we performed repertoire-based BCR selection and corresponding functional characterization against viral proteins. In line with previous observations of antigen specificity within the expanded IgG compartment (Agrafiotis et al., 2023; Neumeier et al., 2021), we observed that a large fraction of expanded IgG clones produced antibodies against influenza-derived NP following infection (∼40% of tested clones). This correlated with the high anti-NP IgG serum titers observed. In contrast, IgA serum antibodies failed to bind to NP, which was also partially reflected in the BM compartment, as the vast majority of highly expanded IgA ASC clones failed to exhibit clear binding to viral proteins (only two exclusively IgA clones showed NP specificity). Further investigation of the expanded IgA ASC compartment revealed signs of polyreactivity against self and non-self antigens, albeit at lower ELISA signal levels compared to influenza antigens.

While our initial objective was to explore the impact of immune imprinting on epitope convergence and cross-reactivity of HA- and NA-specific antibodies, we recombinantly expressed 98 antibodies and none of which exhibited specificity against these proteins. Given that we observed both HA and NA reactivity on a serum level, a deeper selection of antibodies could have revealed antigen specificity, as anti-HA and anti-NA ASCs may be either stochastically distributed within the BM compartment, or the mechanisms by which they populate the BM are distinct to that of NP specific ones. Factors contributing to this lack of expanded anti-HA or -NA IgG clones may include antigen abundance, epitope immunodominance, the nature of antigen presentation or the involvement of other molecular mechanisms that influence antibody binding (Heesters et al., 2013, 2014; W. T. Yewdell et al., 2021). Complex B cell dynamics following immune challenge have previously been reported in the memory compartment, with memory B cells arising from lower-affinity germline cells and failing to bind to antigen (Viant et al., 2020; Wong et al., 2020). Moreover, it is possible that some of these antibodies may be specific to alternative conformations of influenza antigens that would not be revealed by the ELISA assays performed here. GC B cell responses to unknown antigens or non-native forms of immunogen were shown to elicit SHM and clonal expansion following HA immunization (Kuraoka et al., 2016). These “dark antigens” could be related to misfolded proteins or proteolytic degradation products which could be exposed following infection. Finally, the use of monomeric HA for antibody screening may have limited the detection of HA-binding antibodies, whereas employing a trimeric HA form might have yielded additional insights.

Antibodies are largely thought to provide protection in extracellular spaces through Fc-mediated effector functions such as entry-blocking, complement activation and phagocyte recruitment (Chandler et al., 2023). However, many different viruses, including influenza, can also be neutralized inside the cell by antibodies targeting a wide range of viral proteins. This can include proteins exposed on the surface of viral particles or infected cells, or internal proteins only exposed in the cytosol (Bottermann & Caddy, 2022; Chandler et al., 2023; Kok et al., 2023; Mazanec et al., 1995). Given that NP is the most abundant viral protein in infected cells (Y. Hu et al., 2017), in conjunction with the high NP serum titers observed (Sukeno et al., 1979; Vanderven et al., 2022), it seems intuitive that several of the highly expanded ASCs were specific against this viral target. Several studies have shown that anti-NP antibodies can passively transfer protection to naive recipient mice (Carragher et al., 2008; LaMere et al., 2011), however, a similar study demonstrated that human anti-NP antibodies were not protective *in vivo* in a passive transfer mouse model (Vanderven et al., 2022). Unlike external viral antigens, the internal NP is highly conserved (>90%) among all influenza A strains, making it an attractive candidate for vaccination (Y. Hu et al., 2017; LaMere et al., 2011; Ma et al., 2022; Sayedahmed et al., 2024). In fact, several recombinant protein-based vaccine candidates currently in clinical trials include highly conserved NP epitopes (L. Hu et al., 2023). Advances in assays measuring antibody-mediated neutralization inside cells will help shed light on the role of anti-NP antibodies against influenza infection and whether targeting this protein can indeed confer broad protection.

The heavily biased IgA repertoire profiles observed here differ from previous scSeq results of BM ASCs from naive mice (Liu et al., 2023) and presumed background expansion levels of immunized mice, where the vast majority of expanded ASCs were IgM and not antigen specific (Agrafiotis et al., 2023; Neumeier et al., 2021). As this work was performed in identical mice but housed in different animal facilities than the previously reported studies, this suggests that animal husbandry may contribute to experimental variability and have important implications regarding the interpretation of scSeq data. Given their expanded and highly mutated nature, the IgA clones observed here may be of mucosal GC origin (Barone et al., 2011), as intestinal IgA+ ASCs with substantial SHM loads have been previously observed under immune-challenged and homeostatic conditions (Benckert et al., 2011; Lemke et al., 2016; Liu et al., 2023). While the majority of expressed IgA ASCs failed to exhibit clear binding towards viral proteins, some antibodies showed modest change in ELISA signal compared to a negative control antibody. These expanded IgA clones, rather than having high antigen affinity or even influenza specificity, might be driven by different specificity cues taking place in the infected mucosal tissue (Bunker et al., 2017; Ichinohe et al., 2011; Nowosad et al., 2020; Tamura et al., 1990). Previous studies have suggested that influenza induced antibodies were also commonly polyreactive (Andrews et al., 2015; Bajic et al., 2019; Guthmiller et al., 2020). Screening for reactivity against tissue-lysates, microbial components, and other self and non-self antigens suggested reactivity of four antibodies against two or more antigens (20% of expanded IgA clones tested). While we did not detect signs of polyreactivity for the majority of expanded IgAs, it could be that our own methods were not sensitive enough or our antigen panel size was too small to discover the targets of these ASCs. Further studies are required to provide direct evidence of the origin of these IgA clones and understand the role they play during infection or immunization.

In the present study all antibodies were expressed as IgG2c fusions to compare their reactivity independent of their monomeric or polymeric nature. While expressing these antibodies in their native form could have provided additional insights (Bohländer, 2023; Lohse et al., 2011; Okuya et al., 2020; Suzuki et al., 2015), there are still many challenges associated with polymeric IgA expression, mainly because of their complex glycosylation patterns (de Sousa-Pereira & Woof, 2019). In this monomeric IgG form, the specificity assays performed here pertain to the variable antigen-binding site of these antibodies. However, given the polymeric nature of IgA and secretory IgA (SIgA) (Pabst, 2012), a major question still remains as to their non-canonical binding properties (binding via any other constant part of the IgA molecule). In this context, the range of recognized antigens can be broadened by glycan-mediated binding leading to interactions of sufficient strength and valency to drive clearance of infected cells (Mantis & Forbes, 2010; Pabst, 2012; Pabst & Slack, 2020). The possibility of cooperation between different binding modalities of the IgA molecule could play an important role in stabilizing antigen-IgA interactions, however, this has not been extensively studied and functional correlates are unclear.

Collectively, our findings highlight the intricate relationships within the ASC repertoire in homeostasis and following influenza infection. The persistence of highly expanded IgA ASCs following infection, along with transcriptional and repertoire features that appear unaffected by prior immune history, highlights the complexity of mucosal immune responses to respiratory pathogens. The mystery surrounding the role of these IgA clones and their prevalence within the BM compartment even after influenza infection prompts further exploration into potential cues within mucosal tissues. Finally, the identification of expanded influenza-NP specific ASCs highlights the need for further investigation into the role of anti-NP antibodies and their contribution to broad protection against influenza infection.

## Methods

### In vivo animal studies

All animal experiments were performed in accordance with institutional guidelines and Swiss federal regulations. Experiments were approved by the veterinary office of the canton of Zurich. For IAV infection, female 6-8 weeks old C57BL/6 mice (Janvier) were infected with 200 PFU of PR8 (A/PuertoRico/8/1934 (H1N1)) or X31 (a reassortant virus that contains the six internal segments from PR8 and the glycoprotein-encoding segments (HA/NA) from strain A/Aichi/1968 (H3N2)) (LD50 = 1250 PFU) intranasally (i.n.). The X31 stock was egg grown and the PR8 stock was generated and propagated on MDCK cells.

### Isolation of bone marrow B cells

For the BM, the tibia and femur bones were flushed in ice-cold RPMI containing 10% FCS buffer with 10 ng/mL IL6 (216-16, Peprotech). An RBC lysis step was performed in 4 mL ammonium-chloride-potassium lysis buffer for 1 min at room temperature and subsequently inactivated with 26 mL RPMI containing 10% FCS. BM derived single-cell suspensions were stained with the following antibodies (1:200 dilution) CD138-PE, CD4-APC-Cy7, CD8a-APC-Cy7, NK1.1-APCCy7, Ter119-APC-Cy7, B220-APC, CD19-PE-Cy7 for 30 minutes at 4°C. For homologous and heterologous infections, oligonucleotide barcodes conjugated to PE and streptavidin (1:100 dilution) (TotalSeq™-C0951 PE Streptavidin, TotalSeq™-C0952 PE Streptavidin and TotalSeq™-C0953) were added to biotinylated anti-CD138 antibodies to maintain sample identity following pooling of B cells for each experimental condition. For naive mice and infected mice in batch 2, prior to cell sorting, BM ASCs were pre-enriched using a CD138+ plasma cell isolation kit (17627, STEMCELLs EasySet Mouse APC Positive Selection Kit II) following the instructions of the manufacturer and cells were stained with the following antibodies (1:200 dilution) TACI-PE, B220-FITC, CD4-APC-Cy7, CD8a-APC-Cy7, NK1.1-APCCy7, Ter119-APC-Cy7, CD19-PE-Cy7, CD138-APC. Individual mice were stained with oligonucleotide barcode conjugated anti-mouse antibodies (1:100 dillution) (TotalSeq™-C0301, TotalSeq™-C0302 and TotalSeq™-C0303). Cell sorting was performed gating on CD138+ B cells using a FACSAria with FACSDiva software into complete RPMI.

### Single-cell sequencing of antibody repertoires

For single-cell sequencing, antibody secreting B cells (CD138hi) were FACS-sorted from naive and influenza infected mice 15 and 25 days after single and double infections respectively. Single-cell sequencing libraries were constructed from the isolated BM and lung CD138hi B cells following the 10x Genomics’ protocol (CG000330 Rev G). Briefly, single cells were co-encapsulated with gel beads in droplets using 5 lanes of one Chromium Single Cell K Chip (10x Genomics, 1000287) with a target loading of 13,000 cells per reaction. scRNA (GEX) and BCR (V(D)J) library construction was carried out using the Chromium Next GEM Single Cell 5’ Kit v2 (10x Genomics, 1000263), the Chromium 5’ Feature Barcode Kit (10x Genomics, 1000541) and the Chromium Single Cell V(D)J Enrichment Kit, Mouse B Cell (10x Genomics, 1000255) according to the manufacturer’s instructions. Final libraries were pooled and sequenced on the Illumina NextSeq 500 platform (mid output, 300 cycles, paired-end reads) using an input concentration of 1.8 p.m. with 5% PhiX.

### BCR and gene expression data analysis

Raw sequencing files arising from Illumina sequencing lanes were supplied as input to the command line program cellranger (v6.0.0) on a high-performance cluster. Raw reads were aligned to the murine reference genome (mm10) and subsequently supplied into the VDJ_GEX_matrix function of the R package Platypus (v3.1) (Cotet et al., 2023), which relies on the R package Seurat (v4.0.3) (Papalexi & Satija, 2017), and generates an integrated GEX and V(D)J data object. Annotations from GEX were transferred to VDJ and vice versa by matching cellular 10 × barcodes. Cells with more than 15% mitochondrial reads were filtered out and the gene expression was log-normalized with a scaling factor of 10,000. To mitigate batch effects, data integration was performed using Harmony (Korsunsky et al., 2019). Uniform manifold approximation projection (UMAP) was calculated, and feature plots were produced using the FeaturePlot function from Seurat. Dottile plots showing the expression of genes in Seurat clusters were created using the GEX_dottile_plot function from Platypus. Differentially expressed genes were detected using the FindMarkers and FindAllMarkers functions from Seurat and visualized using either the DoHeatmap function from Seurat or the EnhancedVolcano function from the R package EnhancedVolcano (v1.20.0). Cells were assembled into clones based on identical V and J germline gene assignment and a nucleotide similarity in the CDRH3 and CDRL3 of at least 85%. Clonal frequency was determined by counting the number of distinct cell barcodes for each unique clonotype. Those cells in clones supported by only one cell were considered unexpanded clones, whereas those clones supported by two or more cells were considered expanded. Clonal expansion plots were created using the functions VDJ_clonal_expansion and VDJ_clonal_donut function from Platypus. Clone variants were determined as those sequences with the exact same VH and VL nucleotide sequence. Shannon evenness was quantified by using the VDJ_diversity function from Platypus. SHMs were determined using the VDJ_call_MIXCR and VDJ_plot_SHM functions from Platypus. Clonal overlap was calculated based on identical amino acid CDRH3 sequence overlap. Jaccard indices were calculated by quantifying the intersection between two groups divided by the length of the union of the same groups. Clonal overlap was visualized using the pheatmap function from the R package pheatmap (v1.0.12). Germline gene usage was determined by cellranger’s V(D)J alignment to the murine reference. Isotype was determined based on the constant region alignment per cell or for the majority of cells within each clonal family. In the case that the variable region alignment was provided but the isotype was not recovered, the isotype was labeled as ‘‘Unknown’’. ASCs were identified based on the overlay of ASC marker expressions (Cd138 (Sdc1), Taci (Tnfrsf13b), Xbp1, Irf4, Slpi, Cd319 (Slamf7)) on the UMAP and dottile plots of B cell subset assignments (Mathew et al., 2021). The selected ASC barcodes were subsequently supplied into the VDJ_GEX_matrix function from Platypus. For subsequent sequence selection for mAb production, only cells that had one heavy and one light chain were considered. The width of each bar in Figure 4A was normalized based on the total amount of cells in that clone and the total amount of cells per 10x capture.

### Antibody sequence selection and expression

ASCs clones were selected for expression based on their clonal expansion and class-switched profiles. For each selected clone the most frequent unique nucleotide BCR sequence (VH and VL) was selected. The full-length sequences were annotated using MiXCR (v3.0.1) and the AntibodyForest_network_to_pnp function of Platypus. Antibodies were transiently expressed in HEK 293 Expi cells (A14527, Thermo) using the ExpiFectamine 293 Transfection Kit (A14524, Thermo) and the pFUSE2ss vector system (Invitrogen) as previously described (Vazquez-Lombardi et al., 2018).

### Antigen-specific ELISAs for serum and antibody binding

Viral protein specificity was assessed by performing ELISAs against HA1 (11684-V08H, SinoBiological), HA3 (11707-V08H, SinoBiological), NA1 (40196-VNAHC, SinoBiological), NA2 (40199-V07H, SinoBiological), NP (11675-V08B1, SinoBiological) and viral lysate (NAT41630-100, The Native Antigen Company). High-binding ELISA plates (3590, Corning) were coated with the capturing reagent (viral protein or lysate) in PBS at 2 mg/mL overnight at 4 °C. Plates were washed with PBS plus 0.05% Tween and blocked with PBS supplemented with 2% (w/v) milk (A0830, AppliChem) for 2 h at room temperature. Plates were washed and incubated with serially diluted normalized cell culture supernatant (SN) (normalized starting dilution based on ELISA determined supernatant concentration and then 1:4 serial dilutions) or serum (1 to 100 starting dilution and then 1:4 serial dilutions) for 1 h at room temperature. Commercial anti-HA1 (11684-MM05, SinoBiological), anti-HA3 (11056-MM03, SinoBiological), anti-NP (11675-MM03T, SinoBiological) and in-house produced anti-HA1 (sequence taken from (Mathew et al., 2020)) were used as positive controls and a commercial anti-humanTNFR2 antibody (AHR3022, Invitrogen) and in-house anti-hTNFR2 (sequence taken from (Agrafiotis et al., 2023)) and anti-OVA (sequence taken from (Neumeier et al., 2021)), were used as a negative controls. All commercial antibodies were used according to manufacturer’s recommendations. In addition, in-house produced anti-HA1, anti-hTNFR2 and anti-OVA were expressed in both IgG2C and IgA backbones (related to Figure 4B). For detection, an anti-mouse Igk-HRP (AB99617, Abcam) or an anti-mouse Igl-HRP (AB99625, Abcam) was employed at 1:1500 and 1:2000 respectively. For IgA and IgG titer assessment an anti-mouse IgA-HRP (PA1-74397, Invitrogen) and an anti-mouse IgG HRP (A0168, Sigma) were employed according to manufacturer’s recommendations. Binding was quantified using the 1-Step Ultra TMB-ELISA substrate solution (34028, Thermo) and 1M H2SO4 for reaction termination. Absorbance at 450 nm was recorded on an Infinite 200 PRO (Tecan). Polyreactivity ELISAs were carried out as described above. High-binding ELISA plates were coated with Mucin 5AC (abx068104, Abbexa), S100A8/A9 (8916-S8-050, R&D Systems), LPS (tlrl-pglps, InvivoGen), Flagellin (tlrl-stfla, InvivoGen), Mog1-125 (AS-55150-100, Anaspec), hTNFR2 (310-12, Peprotech), Ovalbumin (A5503, Sigma), Calf Thymus DNA (15633019, Thermo) at 10 μg ml−1 in PBS, Fecal bacteria and tissue lysates (lung, brain, colon and small intestine). Fecal bacterial ELISAs were performed as previously described (Nowosad et al., 2020). Briefly, freshly collected feces from cages of naive C57BL/6 mice were macerated with micropestles in 100 μl of ice-cold PBS per 10 mg of feces and vortexed for 5 min. Large debris was removed by spinning at 400g for 5 min at 4 °C. Supernatant containing bacteria was removed and pelleted by centrifugation at 8,000 g. Bacteria were fixed in 0.5% paraformaldehyde for 20 min with continuous rotation to avoid clumping, then washed three times in PBS. High-binding ELISA plates were incubated overnight with bacterial preparations at an optical density at 600 nm (OD600) of 0.35. For tissue lysates, incised tissues were mashed through 70 μm strainers (130-098-462, Miltenyi biotech) and washed with ice-cold PBS. Samples were incubated with RIPA lysis and extraction buffer (89900, Thermo) for 30 min at 4 °C. Following incubation samples were centrifuged at 14,000 g for 15 min at 4 °C and the supernatant was transferred to a new tube. High-binding ELISA plates were incubated overnight with tissue lysates at X in PBS. Following initial blocking with PBS supplemented with 2% (w/v) milk, plates were incubated with goat anti-mouse Igk (AB2338460, Jackson Immunoresearch) to reduce background signal before incubating with mAb SNs. in addition to the previously mentioned controls, the following commercial antibodies were used as positive controls: anti-Flagellin (mabg-flic-2, InvivoGen), anti-DNA (MAB1293, Merck), anti-MOG (MAB5680, Merck) and anti-Actin (A2228, Sigma). All commercial antibodies were used according to manufacturer’s recommendations.

### Quantification and statistical analysis

Heatmaps depicting clonal overlap were created using the R package *pheatmap* (v1.0.12). Heatmaps displaying differential gene expression were produced using the DoHeatmap function in the R package *Seurat* (v4.0.3). Volcano plots were produced using the R package *EnhancedVolcano* (v1.20.0). Experimental overviews were created using BioRender.com. All other plots were produced using Prism v9 (Graphpad). For serum and mAb binding assays, duplicate measurements were performed. For heatmaps depicting absorbance, only the mean value is indicated. All error bars indicate the SE of mean. Any additional statistical details can be found in the figure legends. Figures were assembled and processed in Adobe Illustrator.

## Supporting information

Table S1 antibody information

## Acknowledgments

We acknowledge and thank Dr. Christian Beisel, Mirjam Feldkamp, Elodie Burcklen, and Ina Nissen at the ETH Zurich D-BSSE Genomics Facility Basel for support and assistance. We thank Di Tacchio Mariangela, Cavallini Chiara, and Gumienny Aleksandra from the D-BSSE FACS facility for experimental support. Funding: This work was supported by an SNSF Ambizione Grant (PZ00P3_208734 to A.Y.), an ETH Career Seed Award (A.Y.), and an SNSF Grant (310030_197590 to N.J.).

## Author contributions

A.A., N.J., A.O., S.T.R., and A.Y. contributed to study design. A.A., R.K., C.P., M.V., H.P., L.S., D,S., V.M.A., K.H. and A.G. participated in the material preparation. A.A., M.V. and D.G. contributed to computational analyses and pipelines. All of the authors contributed to the writing of the manuscript.

## Supplementary Figures

**Figure S1.**
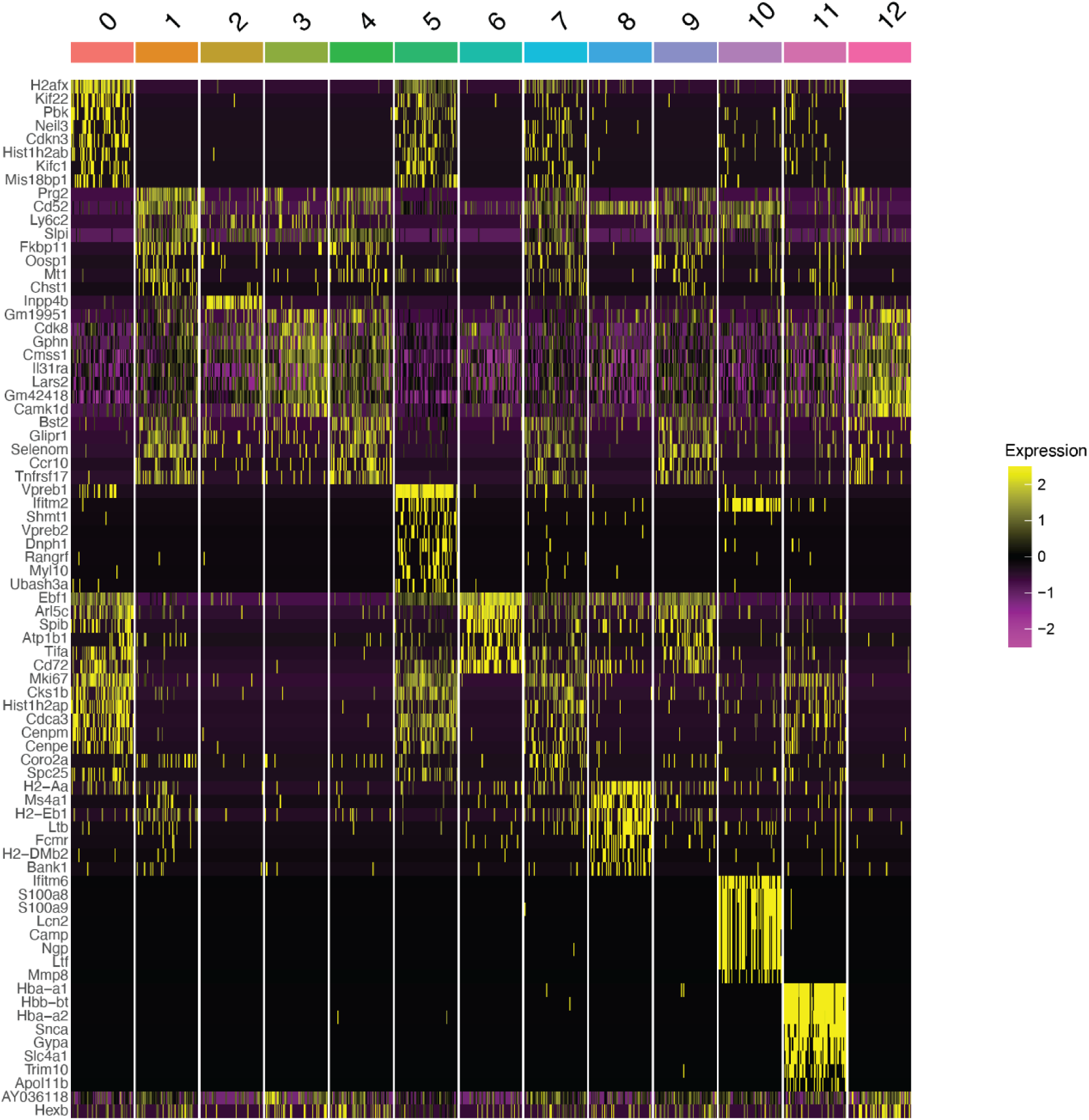
(Related to Figure 1) Heatmap of differentially expressed genes across transcriptional clusters. The order of genes (from top to bottom) corresponds to the highest average log-fold change for each cluster. All genes displayed have adjusted p values < 0.05.

**Figure S2.**
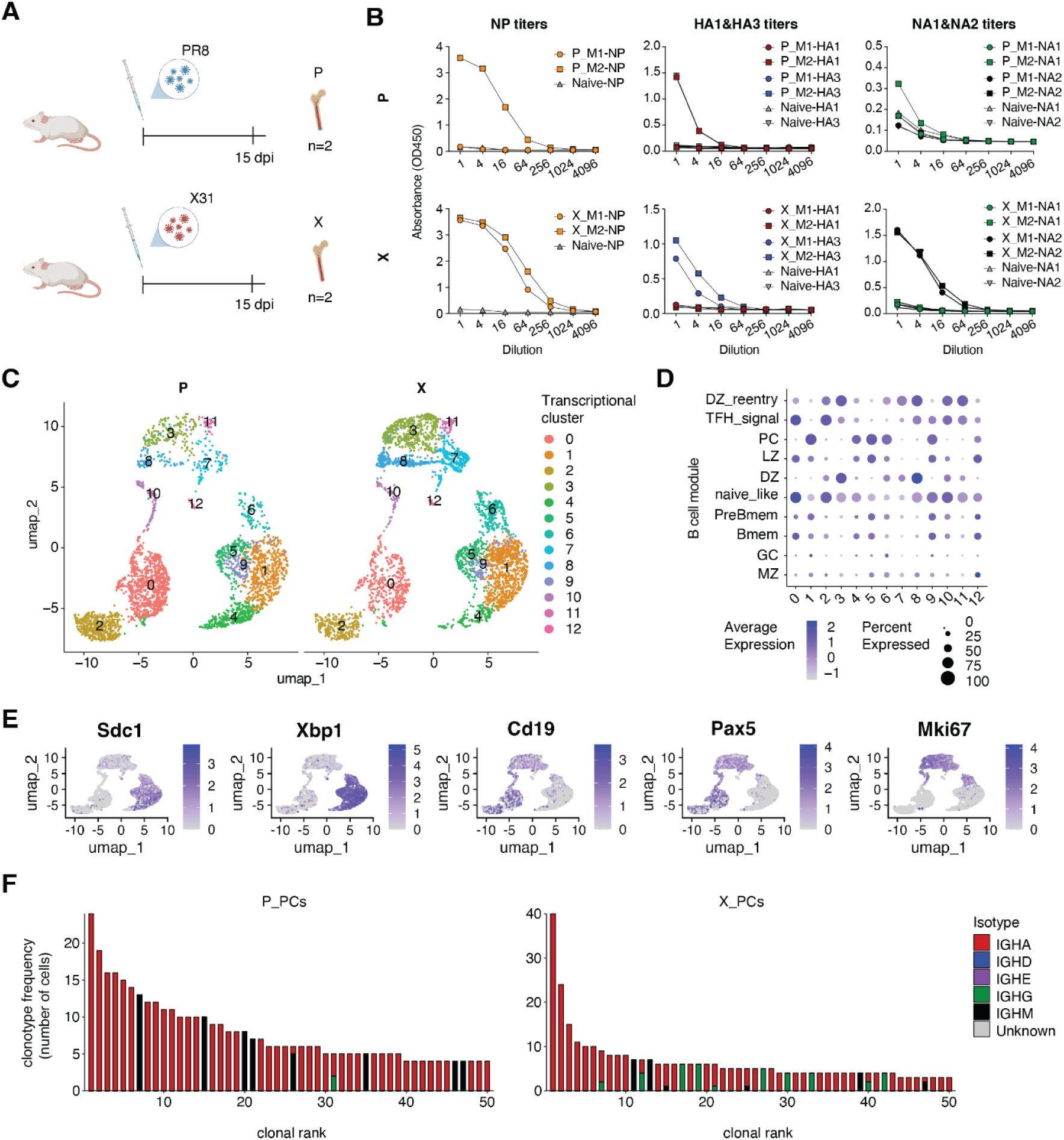
(Related to Figure 2) A) Experimental overview of single influenza infections. B) Per mouse antibody titers measured by ELISA with four-fold pre-diluted serum (1:100) from uninfected mouse (naive) and influenza infected mice (PR8 (P) and X31 (X)) tested against NP (right), HA (middle) and NP (left). Legend indicates condition_batch-capture protein. C) Uniform manifold approximation projection (UMAP)-based total gene expression for lung and BM B cells. D) Dottile plot showing B cell subset assignment across all clusters based on expression of genes defining different B cell phenotypes. The intensity of each dot corresponds to the average expression of all cells within a given cluster, and the size corresponds to the percentage of cells with detectable gene expression. E) UMAP-based gene expression of generic B cell markers. F) Clonal expansion profiles for the 50 most expanded clones colored by isotope (left) and B cell identity (right) per condition and organ.

**Figure S3.**
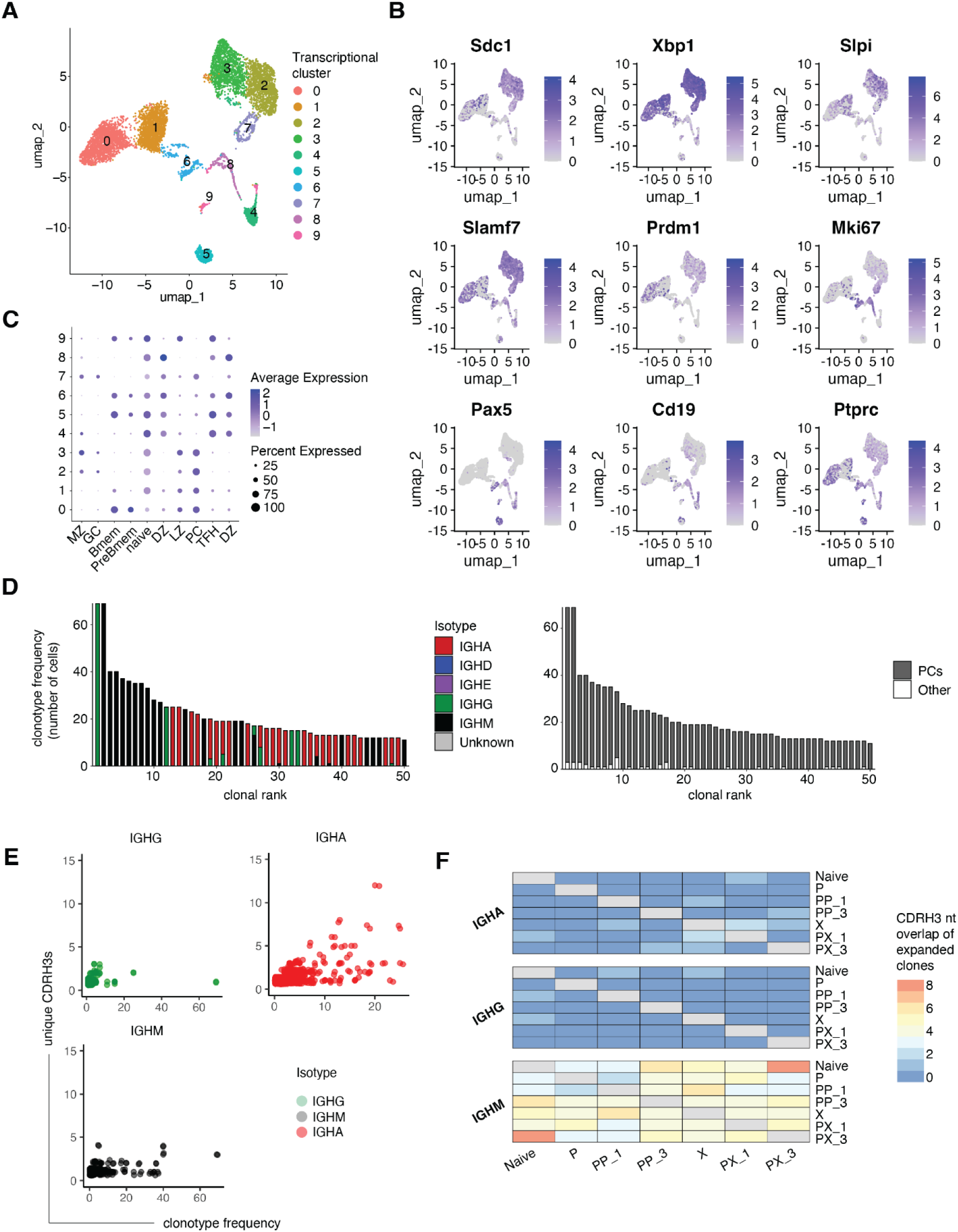
(Related to Figure 2) A) Uniform manifold approximation projection (UMAP)-based total gene expression for BM B cells following scSeq of naive mice (n=3). B) UMAP-based gene expression of generic B cell markers. C) Dottile plot showing B cell subset assignment across all clusters based on expression of genes defining different B cell phenotypes. The intensity of each dot corresponds to the average expression of all cells within a given cluster, and the size corresponds to the percentage of cells with detectable gene expression. D) Clonal expansion profiles for the 50 most expanded clones colored by isotope (left) and B cell identity (right). E) Correlation between the number of unique CDRH3 sequence variants (aa) and the number of cells per clone. F) CDRH3 nt sequence overlap of expanded cells per isotype across conditions.

**Figure S4.**
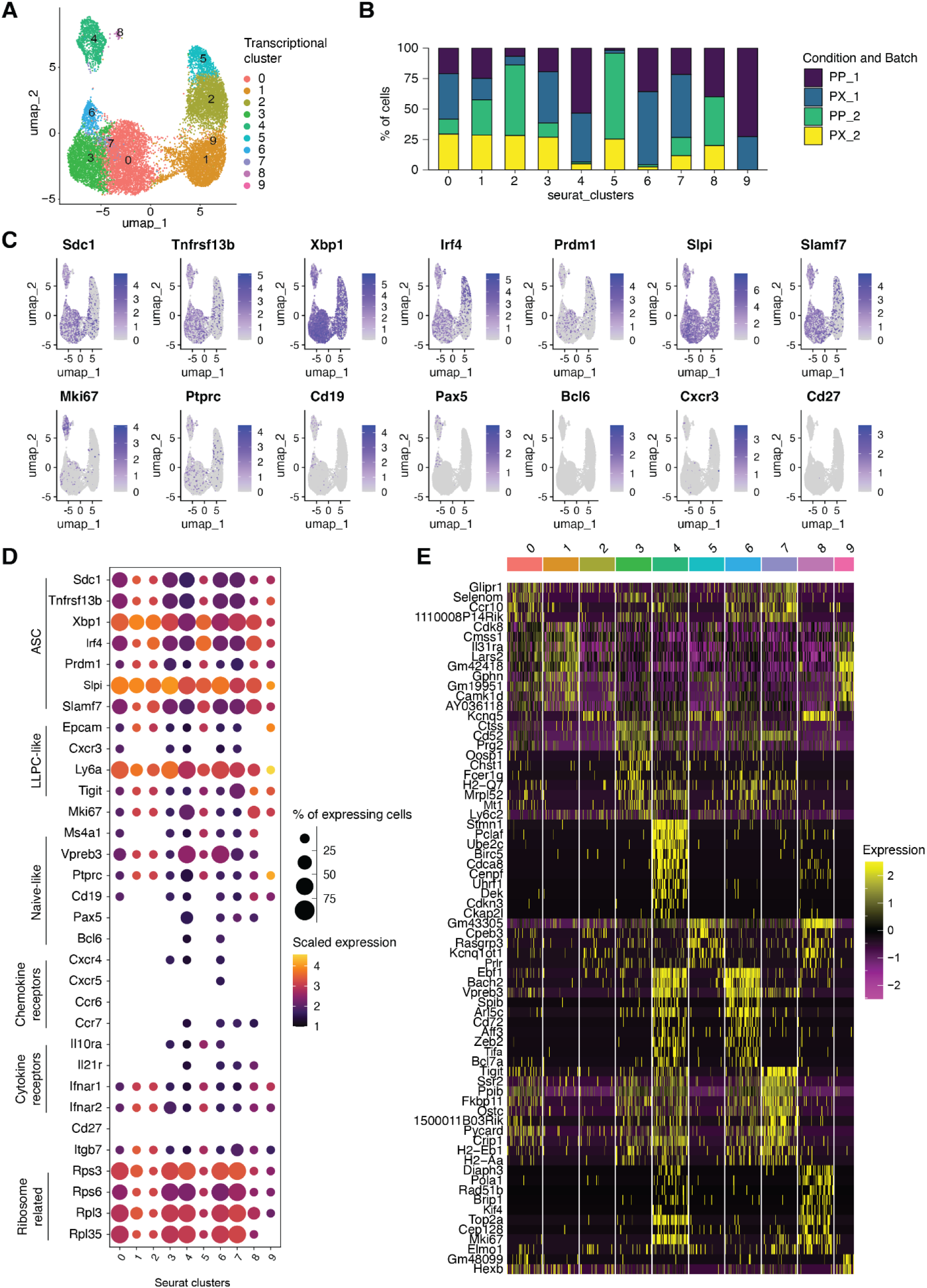
(Related to Figure 3) A) UMAP-based total gene expression for BM ASCs following PP and PX infections. B) Condition and Batch distribution across transcriptional clusters. C) UMAP-based gene expression of generic B cell markers. D) Gene expression levels of B cell markers separated by cluster membership (x-axis). The size of the dot corresponds to the percentage of cells expressing the given marker and the color indicates the mean expression per cell within each cluster. E) Heatmap of differentially expressed genes across transcriptional clusters. The order of genes (from top to bottom) corresponds to the highest average log-fold change for each cluster. All genes displayed have adjusted p values < 0.05.

**Figure S5.**
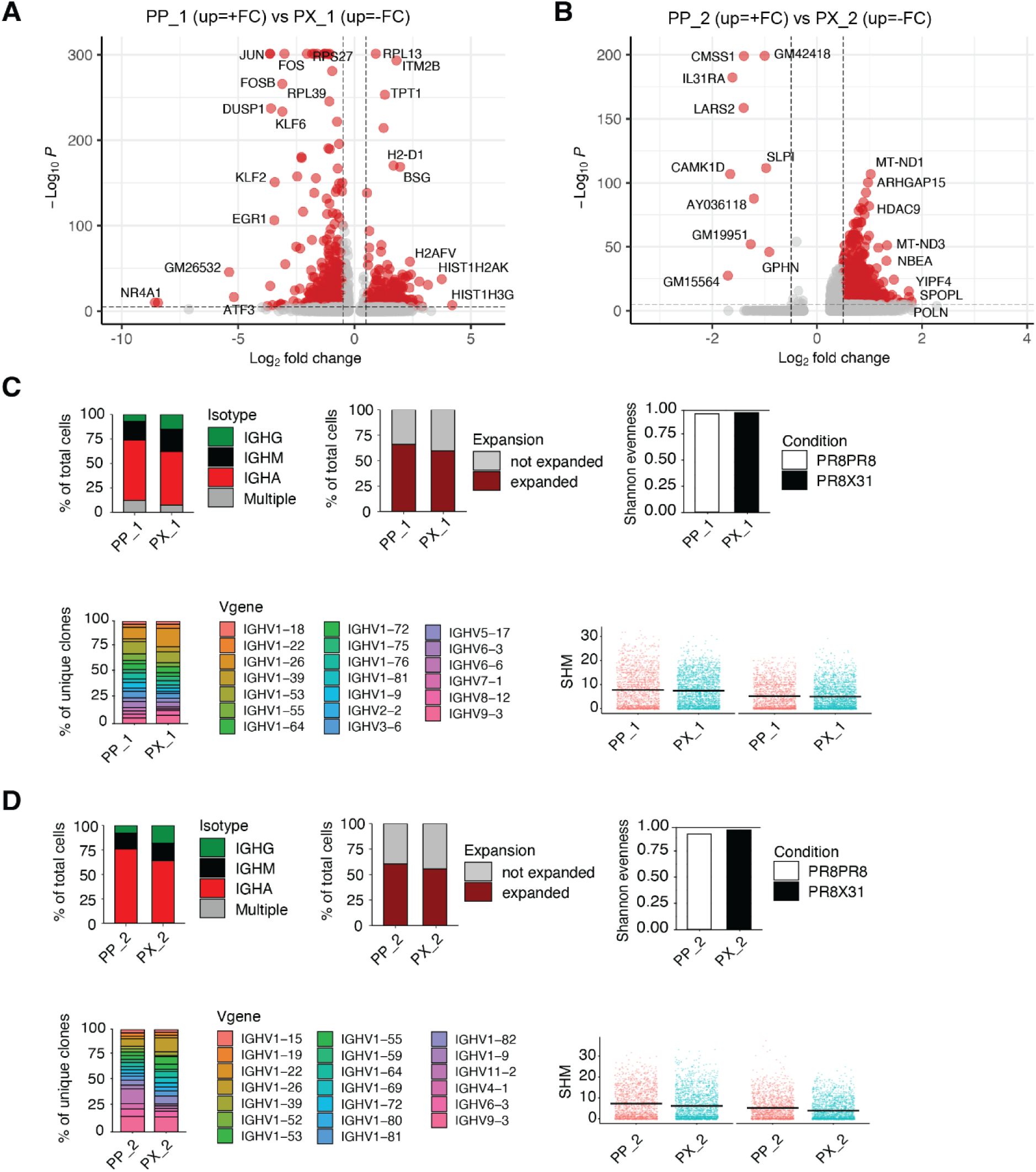
(Related to Figure 3) A-B) Differentially expressed genes between PP and PX ASCs for batch 1 (A) and batch 3 (B). Red dots indicate differentially expressed genes (adjusted P value < 0.01 and average log2 fold change (FC) > 0.5). C-D) Repertoire features (Isotype, Expansion, Shannon evenness, V gene usage and SHMs) for batch 1 (C) and batch 3 (D) derived samples.

**Figure S6.**
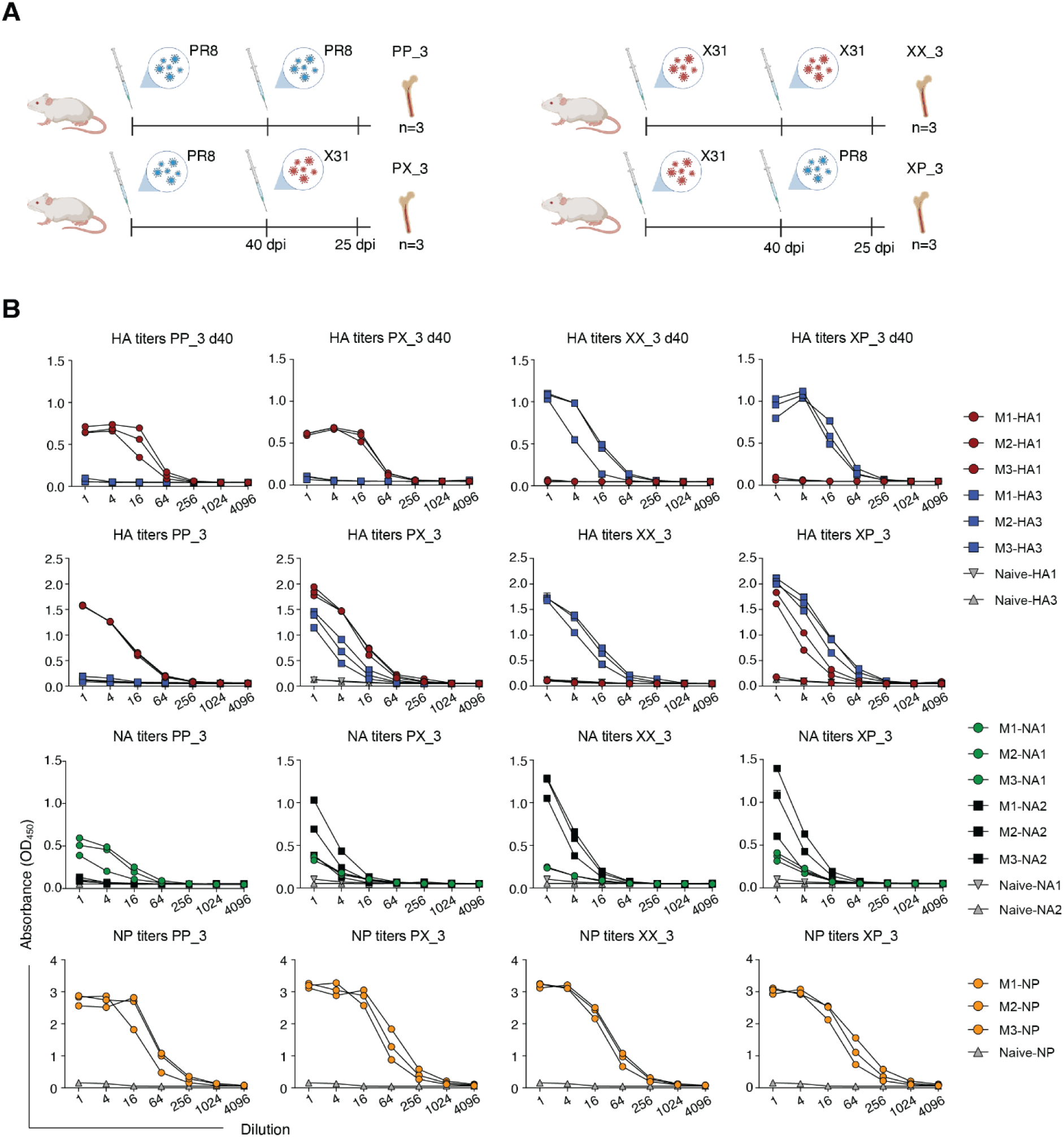
(Related to Figure 3) A) Experimental overview of heterologous and homologous influenza infections with PR8 and X31 strains (batch 3). B) Antibody titers per mouse measured by ELISA with four-fold pre-diluted serum (1:100) from uninfected mouse (naive) and influenza infected mice (PR8PR8 (PP), PR8X31 (PX), X31X31 (XX) and X31PR8 (XP)) tested against HA, NA and NP. Titers against HA were also tested following primary infection (top column). Legend indicates mouse-capture protein.

**Figure S7.**
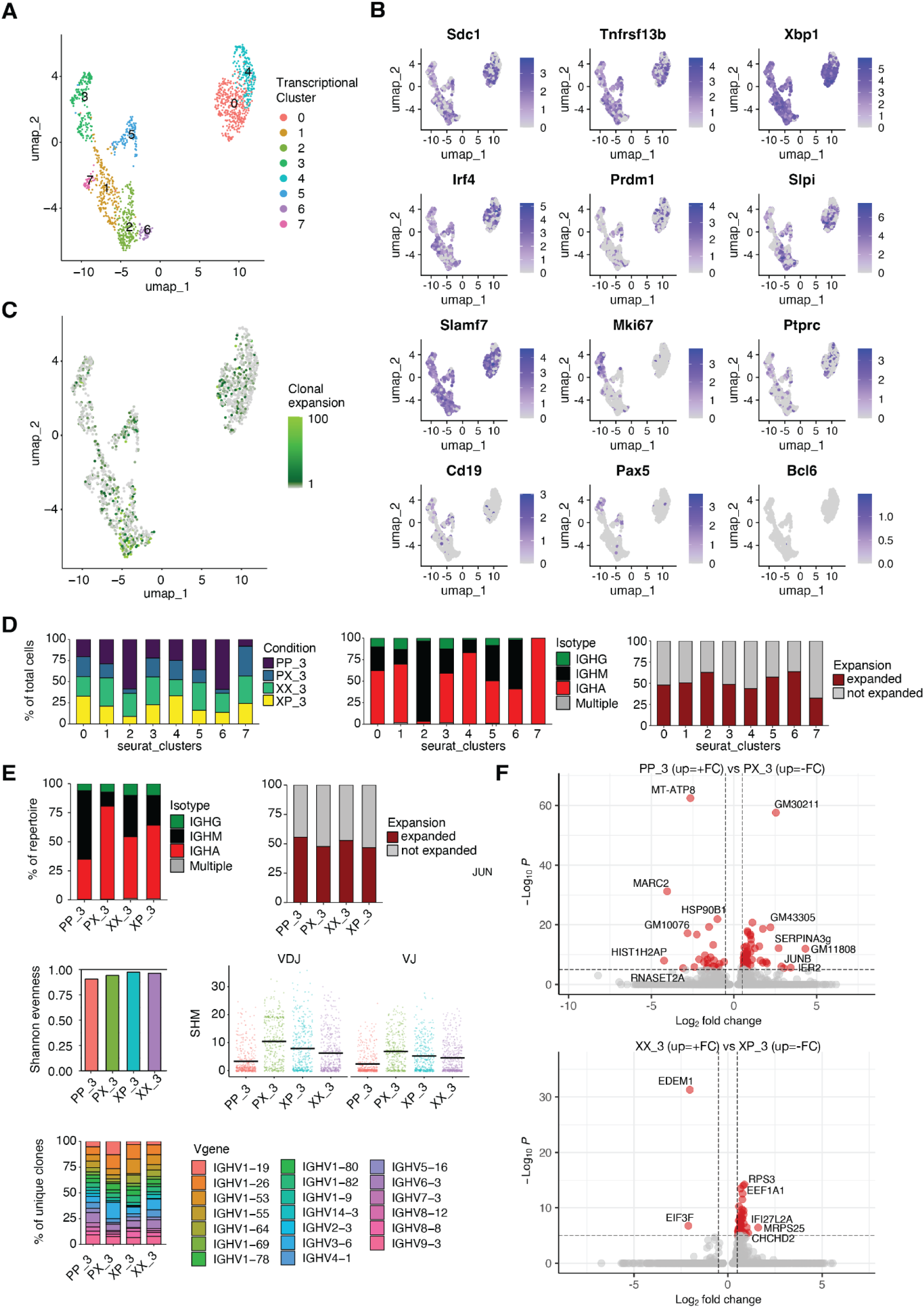
(Related to Figure 3) A) Uniform manifold approximation projection (UMAP)-based total gene expression for BM ASCs following homologous and heterologous infections (batch 3). B) UMAP-based gene expression of generic B cell markers. C) UMAP depicting clonal expansion. Values have been normalized to a scale from 1 to 100, where 1 represents a cell belonging to an unexpanded clone (gray color). D) Condition, isotype and clonal expansion distribution across transcriptional clusters E) Repertoire features (isotype, expansion, Shannon evenness, V gene usage and SHMs) across conditions. F) Differentially expressed genes (adjusted P value < 0.01 and average log2 fold change (FC) > 0.5) between homologous and heterologous samples.

**Figure S8.**
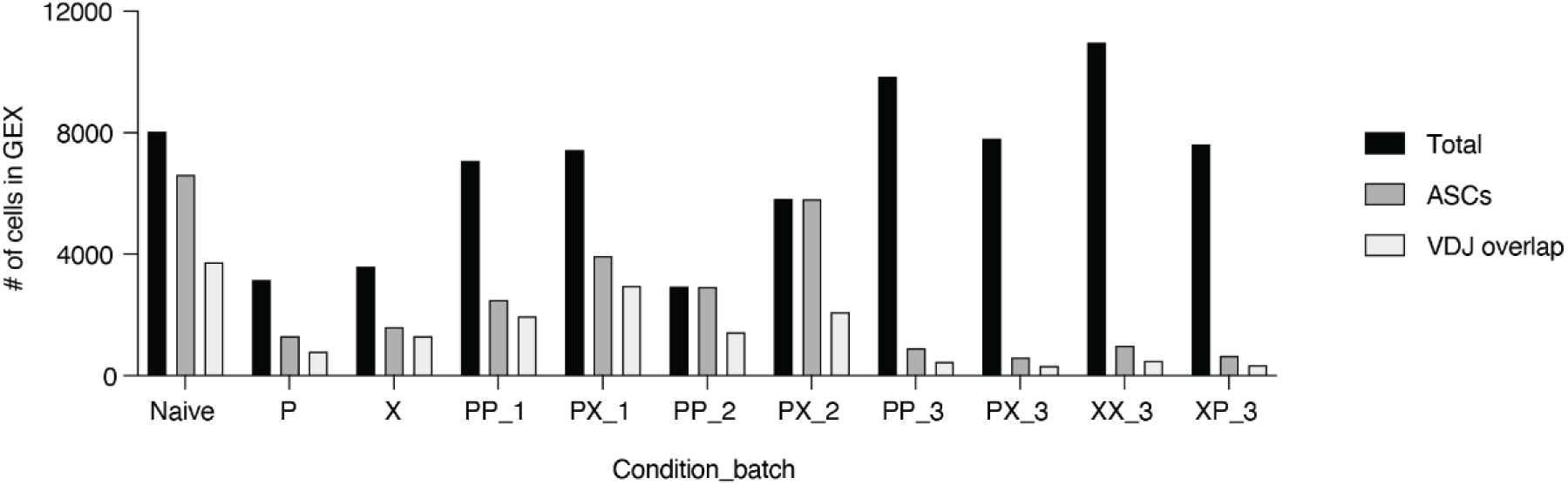
(Related to Figure 3) Number of cells per condition and batch. Black color indicates total number of cells following scRNA sequencing and quality control filtering. Dark gray color indicates number of cells following filtering for ASC markers. Light gray color indicates overlap of ASC barcodes with scBCR cell barcodes.

**Figure S9.**
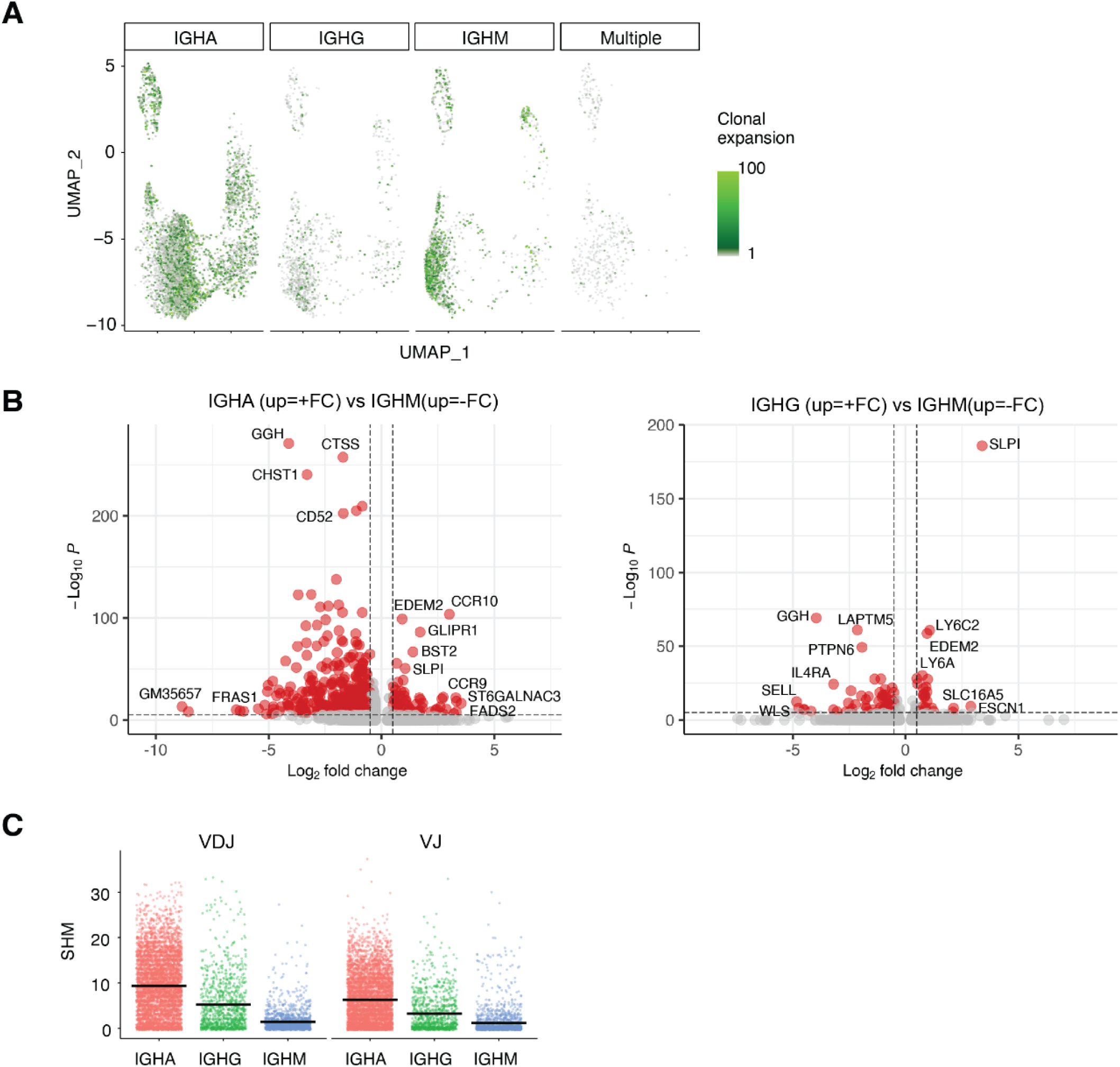
(Related to Figure 3) A) UMAP depicting clonal expansion split by isotype. Values have been normalized to a scale from 1 to 100, where 1 represents a cell belonging to an unexpanded clone (gray color). B) Differentially expressed genes between different isotypes. C) Heavy chain (VDJ) and light chain (VJ) nucleotide somatic hypermutations (SHMs) per isotype.

**Table S1.** (Related to Figure 4) Antibody information for expressed clones (additional csv).

